# Recurrence of *de novo* mutations in families

**DOI:** 10.1101/221259

**Authors:** Hákon Jónsson, Patrick Sulem, Gudny A. Arnadottir, Gunnar Pálsson, Hannes P. Eggertsson, Snaedis Kristmundsdottir, Florian Zink, Birte Kehr, Kristjan E. Hjorleifsson, Brynjar Ö Jensson, Ingileif Jonsdottir, Sigurdur Einar Marelsson, Sigurjon Axel Gudjonsson, Arnaldur Gylfason, Adalbjorg Jonasdottir, Aslaug Jonasdottir, Simon N. Stacey, Olafur Th. Magnusson, Unnur Thorsteinsdottir, Gisli Masson, Augustine Kong, Bjarni V. Halldorsson, Agnar Helgason, Daniel F. Gudbjartsson, Kari Stefansson

## Abstract

*De novo* mutations (DNMs) cause a large fraction of severe rare diseases of childhood. DNMs that occur in early embryos may result in mosaicism of both somatic and germ cells. Such early mutations may be transmitted to more than one offspring and cause recurrence of serious disease. We scanned 1,007 sibling pairs from 251 families and identified 885 DNMs shared by siblings (ssDNMs) at 451 genomic sites. We estimated the probability of DNM recurrence based on presence in the blood of the parent, sharing by other siblings, parent-of-origin, mutation type, and genomic position. We detected 52.1% of ssDNMs in the parental blood. The probability of a DNM being shared goes down by 2.28% per year for paternal DNMs and 1.82% for maternal DNMs. Shared paternal DNMs are more likely to be T>C mutations than maternal ones, but less likely to be C>T mutations. Depending on DNM properties, the probability of recurrence in a younger sibling ranges from 0.013% to 29.6%. We have launched an online DNM recurrence probability calculator, to use in genetic counselling in cases of rare genetic diseases.

## Main text

Humans develop from cell populations derived from a single zygote. The proliferation and differentiation of cells are controlled by a series of developmental programs. One of these, primordial germ cell specification (PGCS), occurs in developmental weeks 2-3 around gastrulation, i.e. before or during the formation of the three major germ layers that yield all of the tissues of the embryo. After the PGCS, the primordial germ cells migrate from the yolk sac to the gonads and proliferate rapidly^1,2^.

Most ssDNMs occur before PGCS or early in the proliferation stage allowing them to become common enough to be transmitted to more than one offspring^3,4^. DNMs mark the progeny of the mutated cell and thus their cell lineage”. Further, ssDNMs can often be found in the somatic cells of parents^8-11^. Therefore, ssDNMs in large sibships are informative about the relationship between cell lineages present in somatic and/or germ cells of the parents.

Standard approaches to search for DNMs in genomic data from parent-offspring trios dismiss candidates with read support for the mutated allele in the parents^12-14^. The rationale is to remove DNM candidates where a parent is in fact heterozygous. Such filters come at the cost of removing DNMs, whose alleles arose in a cell lineage before PGCS^13^.

Despite the fundamental importance of primordial germ cell proliferation and development, little is known about the number of early DNMs and lineages in the population of cells destined for PGCS in humans. To explore this, we sought ssDNMs in 251 parent pairs with two or more offspring (1,007 sibling pairs; Supplementary Table 1; Fig. 1ab). The parental haplotypes present in sibships allow us to find DNMs by identifying genotype discordance between groups of siblings sharing a haplotype (Fig. 1b). We used the ssDNMs to reconstruct cell lineages that gave rise to both somatic cells and germ cells in parents, shedding a light on the DNMs occurring in the earliest stages of the developing embryo.

**Figure 1:**
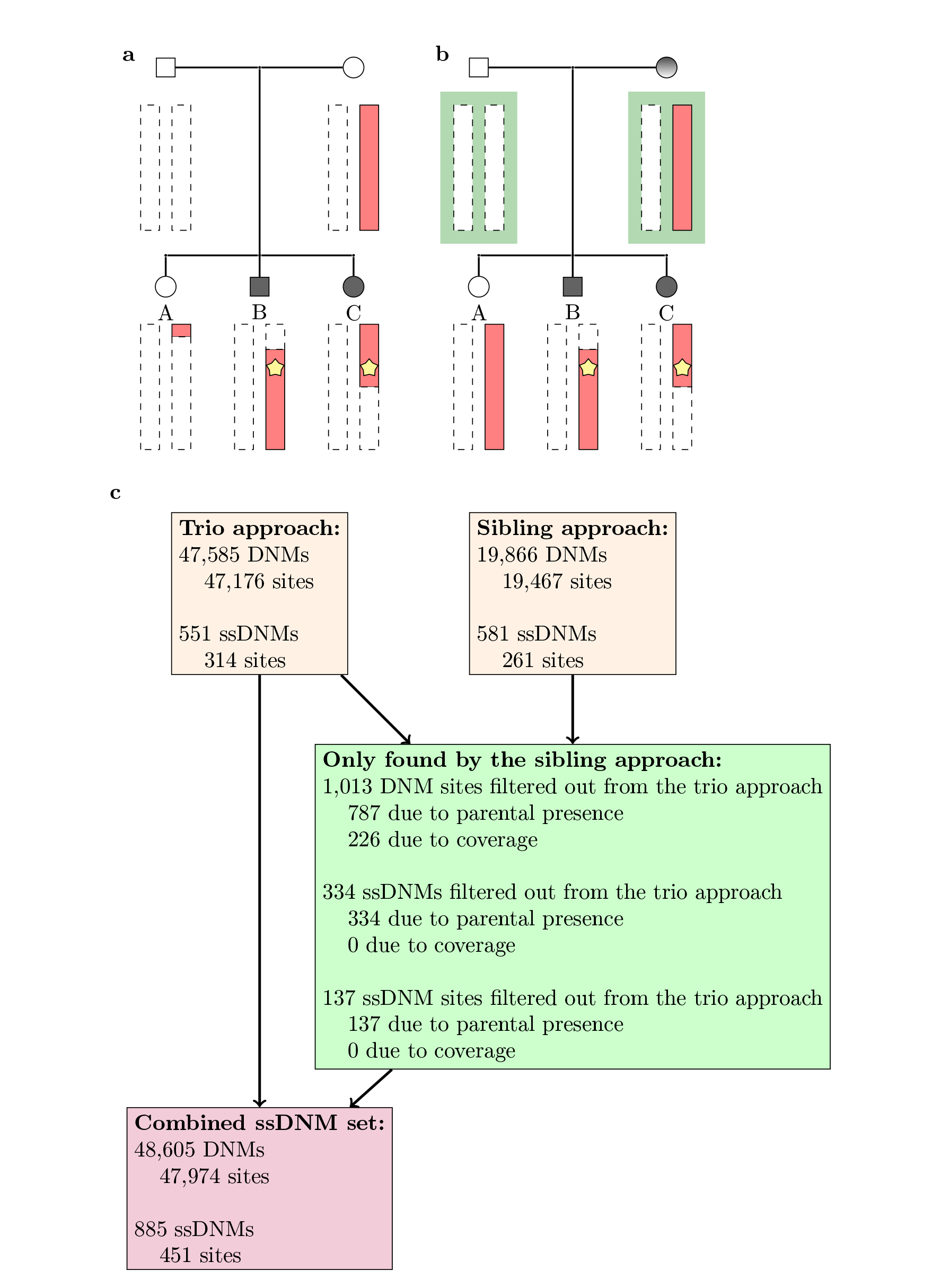
View of the ssDNM extraction. **(a)** ssDNM extraction using genotype comparison to parents. The mutation is determined to be a DNM if the genotypes between the parents and the offspring differ. If there are two or more reads supporting the alternative allele in the parents the mutation is not considered as a DNM candidate, **(b)** The DNM extraction using genotype comparison to siblings. The mutation is determined to be a DNM if the siblings’ genotypes differs despite sharing the same set of parental haplotypes at the locus. The DNM determination is carried out regardless of the genotype of the parents as indicated by the green rectangles, **(c)** The number of DNMs and ssDNMs extracted with each method.

Germline DNMs can be a cause of recurrence of severe genetic diseases among siblings^3^. For example, maternal X chromosome DNMs transmitted to male offspring are a source of recurrence of Duchenne and Becker muscular dystrophies^15,16^, X chromosome DNMs transmitted to female offspring account for recurrence of Rett syndrome^17^, and autosomal DNMs account for recurrence of tuberous sclerosis^18,19^ and osteogenesis imperfecta^20^.

When a DNM causes a disease, it is important to provide the parents of the affected child with a probability of recurrence in future children. Whereas a recessive pattern of inheritance is associated with a recurrence probability of 25% and a dominant one with 50%; current estimates of recurrence of a DNM among siblings range from 0.048% to 9.4% depending on the parent-of-origin, older sibling carrier status and presence of somatic mosaicism^10^. However, these estimates are based on a theoretical model of the propagation of cells of the developing human embryo and are therefore not suited for clinical applications. DNMs are rare and DNM sharing even rarer, therefore the estimation of recurrence probabilities requires a large amount of high quality data. By characterizing the impact of parent-of-origin, read frequency in parent, older sibling carrier status, genomic location and mutation class of ssDNMs, we constructed a calculatorio assess the probability of DNM recurrence for use in genetic counselling.

## Results

We searched for ssDNMs in 251 couples with two or more offspring in our previously described set of 1,548 Icelandic trios (whole-genome sequenced to average coverage of 35X coverage)^21^. This set was enriched for large sibships: wherein 13 couples had 8 or more offspring and the largest family had 17 offspring. This resulted in 1,007 pairs of siblings (Supplementary Table 1), where more than half of the pairs (518) come from the large sibships. In order to estimate the levels of somatic mosaicism in the parents more accurately, we whole-genome sequenced the 26 parents of the large sibships to average coverage of 172X. For most of the sibling pairs (966 out of 1,007) parental DNA was extracted from blood, and for the deep-sequenced parents of the large sibships all of the parental DNA was extracted from blood.

We identified 47,585 autosomal DNMs (47,176 sites) in the 251 families by the trio approach (Fig. 1a), i.e. requiring the DNM absence from the parents. Of these we found 551 ssDNMs (at 314 sites). We extended the set of early DNMs by considering mutations that were present in at least one offspring, but not all siblings sharing a haplotype at the chromosomal position of the mutation, but not necessarily absent from the soma (blood and buccal) of the parents (Fig. 1b; Methods). We found 19,886 autosomal DNMs (19,467 sites) and 581 ssDNMs (261 sites) with this approach. We compared the overlap between the two DNM sets (Fig. 1c) and found 1,013 DNM sites that had been filtered out in our previous analysis, whereof 787 were omitted due to reads in parents and 226 due to insufficient coverage in the parents. This parental presence filter is pronounced for ssDNMs, as 334 (137 sites) of the 581 ssDNMs (261 sites) identified by the sibling approach had been omitted due to reads found in parents (57.5%). Thus, while the standard approach of requiring the absence of reads with mutated alleles from parents when identifying DNMs in children leads to only a small fraction of all true DNM sites being omitted (4%; 787/19,467), this is a substantial fraction of ssDNMs sites (52.2%; 137/261). We collated these two data sets of DNMs resulting in a set of 885 ssDNMs at 451 sites (Fig. 1c; Supplementary Table 1).

We used haplotype sharing between siblings in families with several ssDNM sites to reconstruct the phylogenetic relationship of the germ cell lineages underlying the sibling zygotes (Methods; Fig. 2; Supplementary Fig. 1-2). Fig. 2 summarizes the pattern of paternal and maternal ssDNMs among the siblings of the two largest families. The paternal germline tree in the family with 17 offspring consists of two major lineages (17-P-1 and 17-P-2), each comprised of 8 individuals and defined by 9 and 7 ssDNM sites, respectively. Interestingly, one of the offspring (born in 1963) does not share a DNM with any of its 16 siblings. We calculated the fraction of reads supporting the alternative allele (Allelic balance, AB) and found the AB in the blood of the father was up to 8.8% and 47.7% for lineages 17-P-1 and 17-P-2, respectively (Fig. 2a). The ssDNMs with somatic presence in the father are not shared by the different cell lineages (17-P-1 and 17-P-2). We also observed two major lineages (17-M-1 and 17-M-2) for the maternal germline. The maternal soma cells were almost entirely from the 17-M-2 lineage (Fig. 2b).

**Figure 2:**
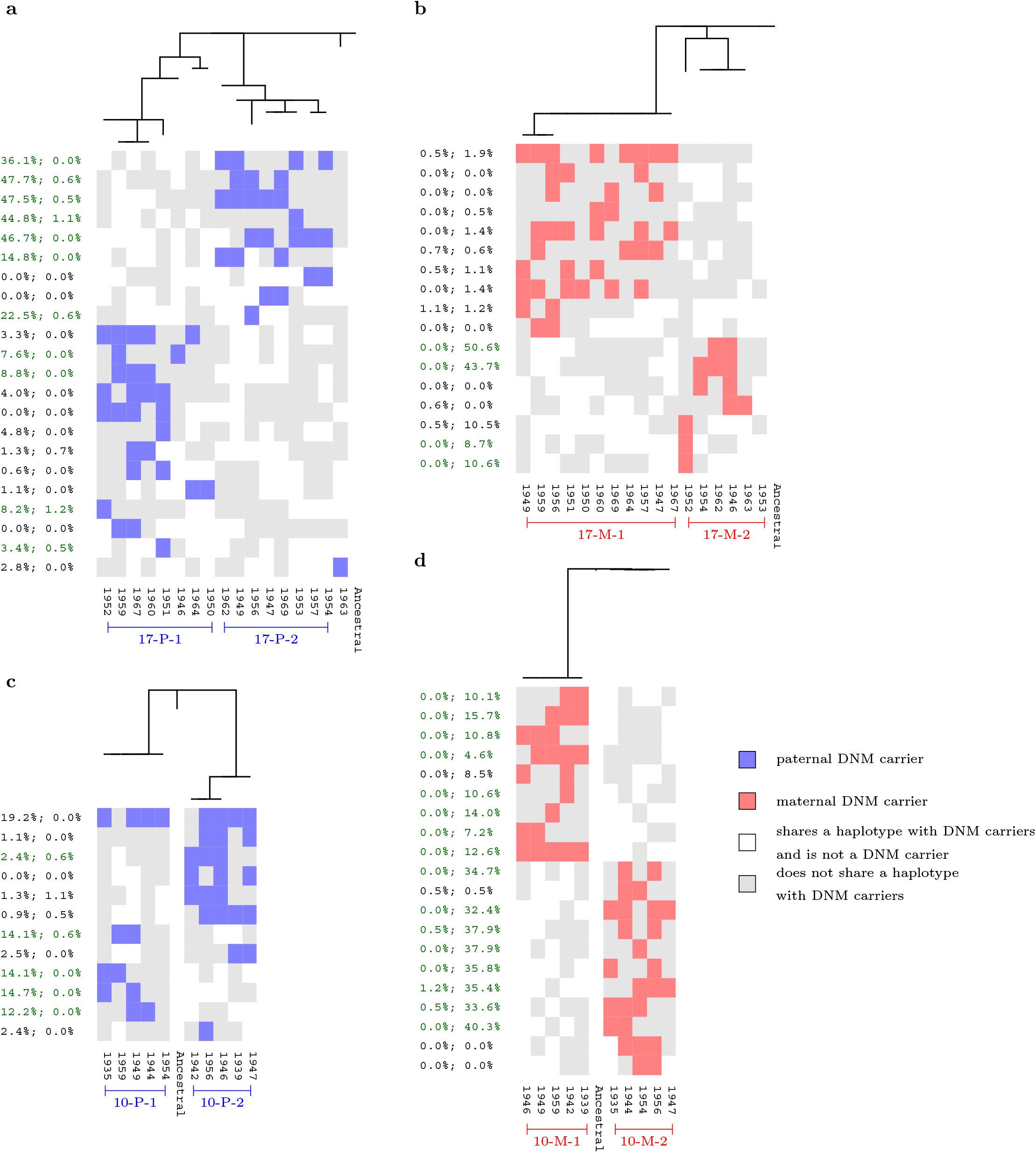
Phylogenetic trees for the germline lineages. **(a)** and **(b)** the male and female germ cell lineages for the 17 offspring family, **(c)** and **(d)** the paternal and maternal germ cell linages for the 10 offspring family. Black and green text correspond to DNMs determined by trio and sibling approach, respectively. Percentages are paternal and maternal ABs, respectively. In addition to ssDNMs, we incorporated DNMs that are somatic mosaic in the parent (≥4 reads supporting the alternative allele in the parent). DNM sites are ordered according to carrier frequency.

The paternal ssDN Ms in the 10 sibling family also form two lineages (10-P-1 and 10-P-2) that differ considerably in the levels of paternal somatic mosaicism (Fig. 2c, AB=12.2%-14.7% vs AB=0.9%-2.5%). A single paternally derived ss DNM (chrl0:32,577,095 G>A) was shared by eight siblings and was present in both major lineages, 10-P-1 and 10-P-2, as well as being present in a mosaic form in the blood of the father (AB=19.2%). In the maternal germline of the 10 offspring family, there were two lineages (10-M-1 and 10-M-2), both with high levels of somatic mosaicism (Fig. 2d, AB=4.6%-15.7% and AB=0-40.3%, respectively).

The mutations in the germline trees shared by several siblings and showing high levels in the parental blood most likely occurred before the PGCS. In contrast, the mutations with no somatic presence and shared by few siblings in the germline trees are consistent with two scenarios, either they are present in the blood at a lower frequency than can be detected in our study or appeared in the germline after the PGCS.

To quantify the relationship between DNM recurrence and parental somatic mosaicism, we compared the ssDNM fraction to the levels of somatic mosaicism of the parents. We found a strong correspondence between the allelic balance in the parents and the ssDNM status (0.127 AB difference; p=1.0·10^-12^; Fig. 3ab and Supplementary Fig. 3). For the 26 deep-sequenced parents (172X average coverage), the sharing by siblings increased from 0.4% when no parental read contained the alternative allele to 13% when 3 or more reads contained the alternative allele. We also identified pairs of DNM sites from the same parent which we could date by the DNM sharing (Fig. 3c), and compared their relative age determined by the sibling sharing to their allelic frequency in the parent. More specifically, if two siblings S1 and S2 share DN M A, but DN M B from the same parent is only observed in sibling S1 even though the siblings share a haplotype from the parent at the position of B, we can conclude that DNM A is older than DNM B. We dated 14,259 pairs of autosomal DNM sites, 11,633 from fathers and 2,626 from mothers. We found that older DNMs are present at greater frequencies in the parent's blood than younger DNMs (p=3.2·10^-2^ and p=1.6·10^-13^ for paternal and maternal DNMs, respectively; Fig. 3de). This supports the hypothesis that older DNMs as determined through the sibling sharing occur more frequently before the PGCS than younger DNMs.

**Figure 3:**
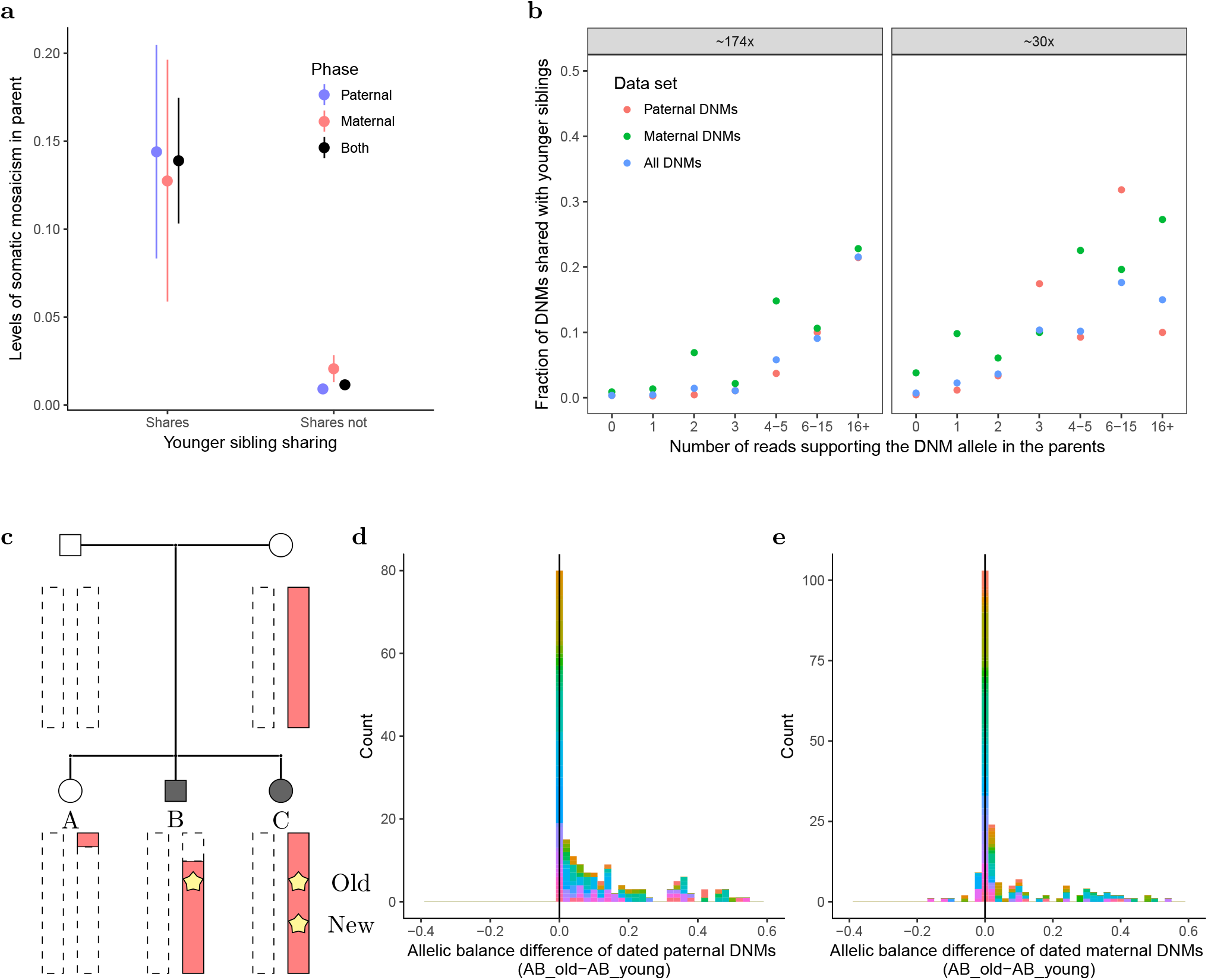
The relationship between germline and somatic mosaicism. **(a)** ssDNM rate against allelic balance in parents. Restricting to the deep-sequenced parents. **(b)** ssDNM rate against total number of reads supporting the alternative allele in the parents. We used DNMs from the trio and sibling DNM approach for **(a)** and **(b). (c)** schematic view of pairwise dating of DNMs. **(d)** the levels of somatic mosaicism for pairwise dated paternal DNMs. **(e)** the levels of somatic mosaicism for pairwise dated maternal DNMs. In **(d)** and **(e)** each family has a different color.

We next compared mutation rates and spectra of paternal ssDNMs to maternal ssDNMs. We previously determined the parent-of-origin of 42,961 DNMs and estimated that 79.4% come from fathers^21^. In contrast, we estimate that only 386 ssDNMs (187 DNM sites; 47.8%) of the 807 (383 DNM sites) phased ssDNMs are of paternal origin. Accordingly, we estimate the paternal and maternal autosomal ssDNM rates to be 1.06% (95%-CI:0.91-1.20%) and 4.52% (95%-CI:3.85-5.18%), respectively. A greater maternal fraction of ssDNMs is expected, since DNMs increase more rapidly with paternal age than maternal age, therefore a greater fraction of maternal DNMs are likely to have occurred early.

A driver mutation can induce proliferation of the mutated germ cell^22^, resulting in its overrepresentation among sperm, which could lead to an increase of ssDNMs with paternal age. This is not the case in our data set. Due to the overall age increase in paternally transmitted DNMs, the fraction of paternal ssDNMs decreases with father's age (decrease of 2.28% per year, 95%-CI:1.94-2.61%; Fig. 4a). The absolute number of paternal ssDNMs remains constant with father's age (3.8·10^-3^ ssDNMs per year, 95%-CI:-4.1·10^-3^-1.2·10^-2^). This indicates that the germ cell diversity of fathers is typically maintained throughout adult life. We see a similar decrease in the fraction of maternal ssDNMs with maternal age (decrease of 1.82% per year, 95%-CI:1.10-2.53%; Fig. 4b). Although the maternal age effect on the number of transmitted DNMs is substantially weaker than the paternal age effect^21^ (0.37 maternal DNMs per year vs 1.51 paternal DNMs per year), the relative increase in the number of DNMs with parental age is similar (4.1% and 3.4% per year for paternal and maternal DNMs for 20 year old parents, respectively).

**Figure 4:**
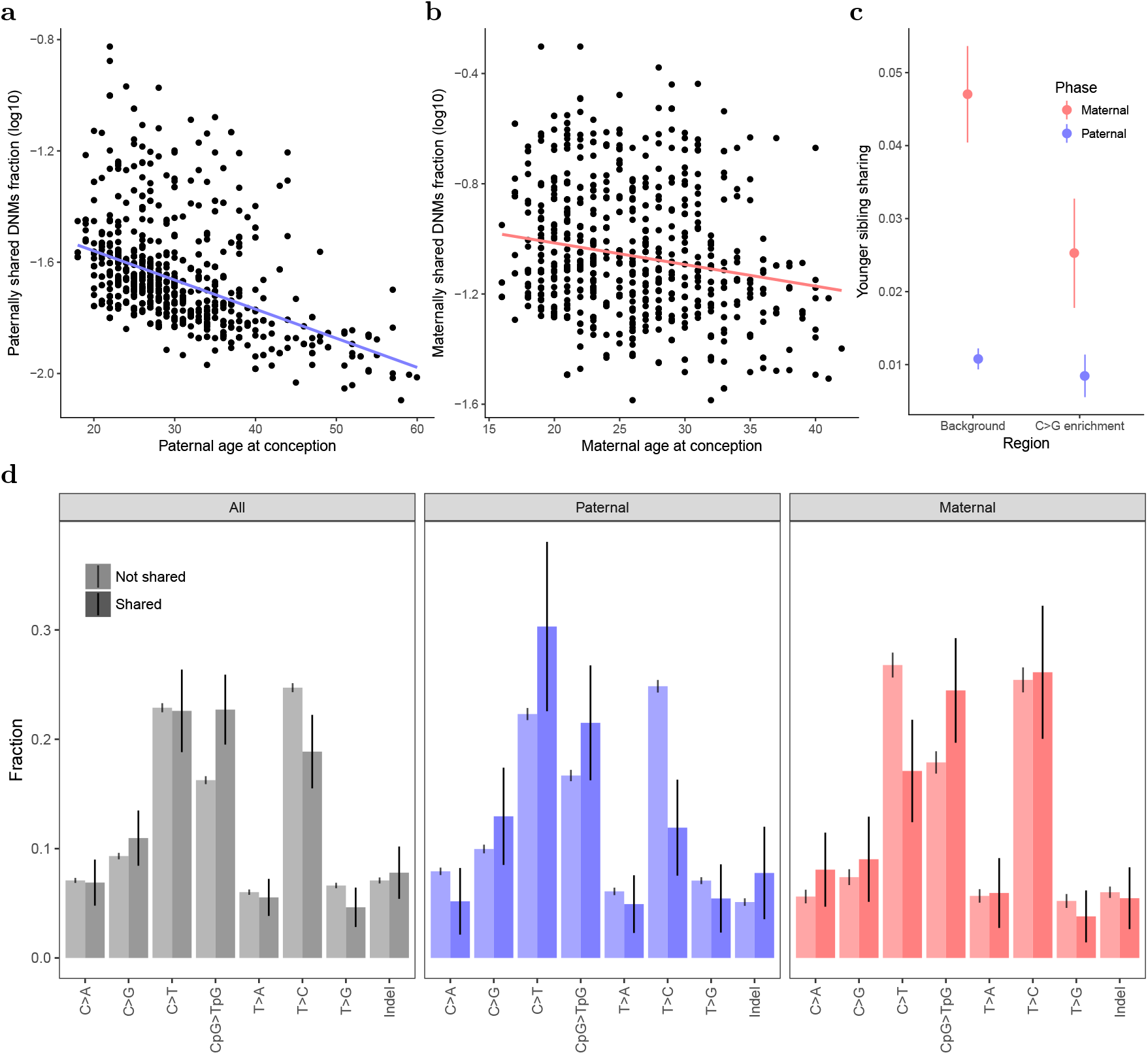
Summary of the shared DNMs from the trio approach. **(a)** maternal ssDNM fraction against paternal age at conception of older sibling, **(b)** paternal ssDNM fraction against maternal age at conception of older sibling. In **(a)** and **(b)** the number of phased ssDNMs was divided by the imputed number of phased DNM pairs and only DNMs from the trio approach were used. A pseudo count of 1 was added to the numerator and denominator of the ratio, **(c)** Fraction of DNM site sharing stratified by region, **(d)** Mutation spectra for DNMs and ssDNMs.

We have shown that the age related DNMs from mothers accumulate in specific regions which are characterized by a high fraction of C>G variants (C>G enriched regions)^21^. Therefore, maternal DNMs within C>G enriched regions should be less likely to be shared by siblings. Indeed, we find a lower maternal ssDNMs fraction within the 10% of the genome most enriched for C>G variants (2.53%, 95%-CI:1.78-3.27%) than in the rest of the genome (4.70%, 95%-CI:4.04-5.36%) (p=5.8·10^-6^; Fig. 4c).

Comparing shared to non-shared DNMs, we tested for enrichment of eight different mutational classes^21^, C>A, C>G, >>T, CpG>TpG, T>A, T>C, T>G and Indel. This revealed a greater number of CpG>TpG transitions (odds ratio=1.52; 95%-CI:1.31-1.76) and fewer of T>C mutations (odds ratio=0.71; 95%-CI:0.59-0.86) among the ssDNMs compared to the non-shared DNMs (Fig. 4d). Furthermore, for ssDNMs there is a higher contribution of T>C (odds ratio=2.61; 95%-CI:1.66-4.10) and lower contribution of C>T (0.47; 95%:0.29-0.75) in the maternal germline.

Our results indicate that the mechanism behind DNMs occurring before and after the PGCS differ. We show how sex differences in germ cell development induce considerable parent-of-origin effects on the spectrum, rates and genomic location (C>G enriched regions) of ssDNMs. These easily accessible attributes of DNMs are currently not utilized in the genetic counselling of DNM recurrence.

The number of DNM transmissions in sibships is informative about the frequency of the DNM in the germline, thus also about the risk of DNM recurrence. More specifically, if two offspring share a DNM, the estimated probability of a transmission to a third offspring increases to 22.6% (95%-CI:20.0%-25.2%) from 1.1% (95%-CI:0.96-1.21%). The increased probability is similarly high for DNMs of paternal and maternal origin, i.e. 20.9% (95%-CI:16.7%-25.0%) and 26.1% (95%-CI:23.3%-29.0%), respectively. This increase from both parents is substantially greater than the estimates from the theoretical modeling of gametogenesis^10^: when parental somatic mosaicism is unknown then 0.096%-0.468% of DNMs from fathers and 4.95% from mothers are ssDNMs, and when parental somatic mosaicism is present in the parent 9.40% from mothers and fathers.

We have launched an online recurrence calculator (http://de-novo-risk.decode.is/UsentesterPassword:tester.pass!) that incorporates the parent-of-origin, genomic position, occurrence in an older sibling, parental age, presence, and levels of parental mosaicism as covariates. Accurate quantification of mosaicism in parents, older sibling information and determination of the DNM phase is of utmost importance for recurrence estimation. However, accurate determination of DNM phase and levels of somatic mosaicism can be impractical for clinicians. We therefore predict phase of the DNM and incorporate the uncertainty of the somatic mosaicism determination of the parents into the calculator by estimating the posterior probability that a parent is mosaic (PPPM; Methods; Fig. 5a). The estimated recurrence probability of a mutation that is present in the blood of a parent (PPPM=1) will be transmitted to a subsequent offspring are 14.7% and 21.4% for fathers and mothers, respectively. Further, the estimated recurrence probabilities are 0.3% and 1.6-1.9% for 30 year old parents if the DNM is not present in the blood of a parent (PPPM=0). In the extreme case, if the older sibling is a carrier and the mother is somatic mosaic the resulting predicted probability is 29.6%. On the other end, if the older sibling is not a carrier, both of the parents are old at conception (father is 60 and mother is 40), the DNM is paternal, and there is no somatic presence in the father then the predicted probability is 0.013%. The range of our directly estimated recurrence probabilities (0.013%-29.6%) is substantially greater than those estimated with a modeling of the mutagenesis in the developing embryo (0.048%-9.4%)^10^.

**Figure 5:**
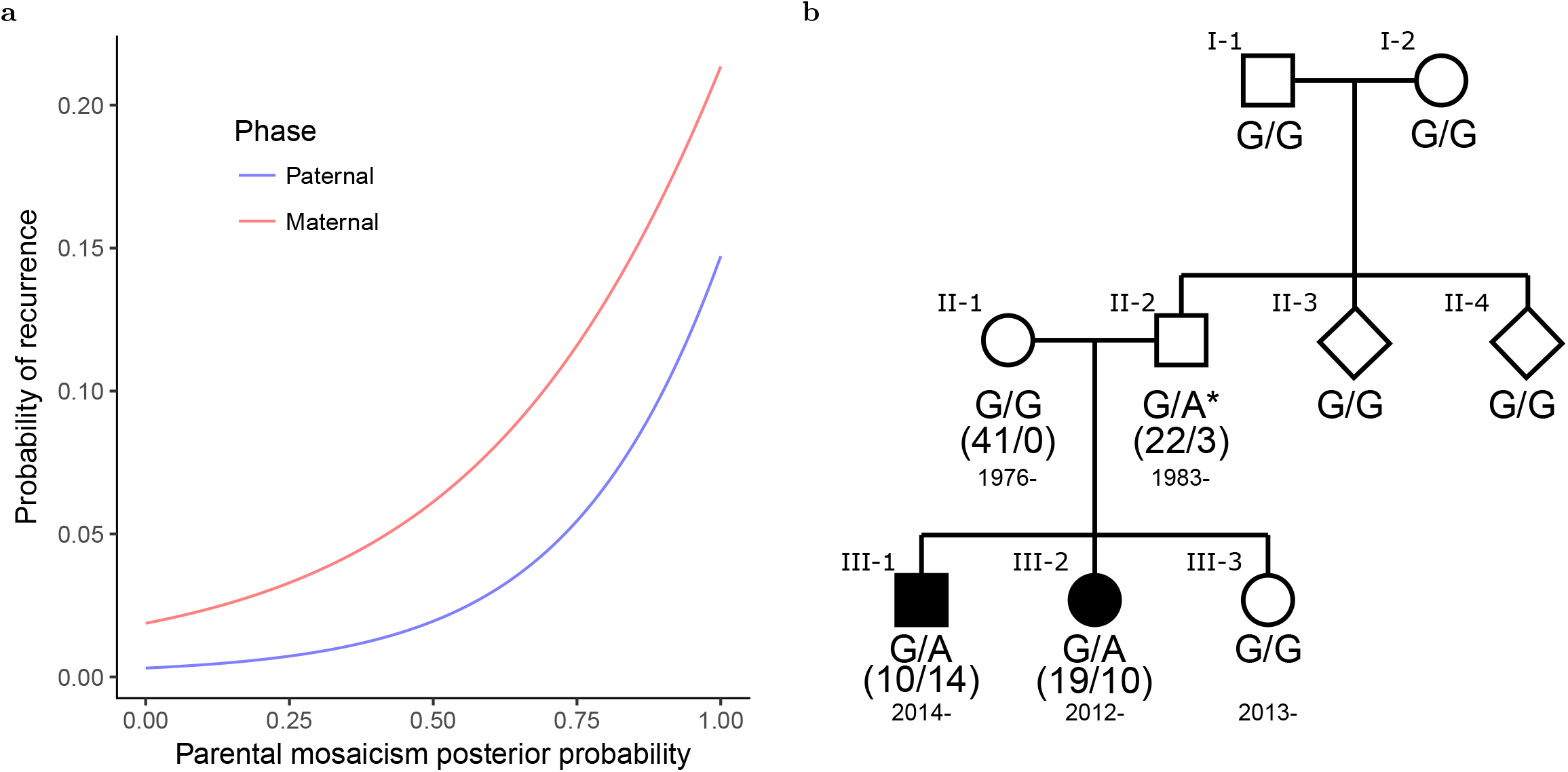
Germline mosaicism in the context of clinical genetics. **(a)** the predicted recurrence probability against the parental mosaic posterior probabilities. For the paternal prediction the maternal somatic mosaic posterior probability was set to 0 while the paternal one was varied across the range [0; 1]. For the maternal prediction the reciprocal was done. The recurrence probability prediction is based on: the father and mother are 30 year old at conception of the proband, there is no information about older siblings, the DNM is phased and not in a C>G enriched region, **(b)** the STXBP1 clinical case. The affection status of the siblings is indicated by blackened symbols. The genotypes at variant chr9:127,682,486 G>A are denoted below the individuals. The mosaic genotype of the father of the affected siblings is marked with an asterisk. The number of reads supporting the reference and alternative allele are depicted in the parenthesis below the genotype.

We were presented with a pair of young siblings not in our set of 1,548 trios with epileptic encephalopathy and their unaffected parents and siblings (Methods; Fig. 5b). We whole-genome sequenced the family and detected a missense mutation p.Gly543Glu (N P_003156.1) of paternal origin in the known epileptic encephalopathy gene, 5TXÖPI^23,24^, shared by the two affected siblings (Methods). In addition to two transmissions of the DNM we found that the DNM is mosaic in the blood of the father (AB=12%), which dramatically changes the genetic counselling for subsequent pregnancies, as the estimated probability of DNM recurrence increases from 0.27% to 22.5% for future offspring of the father.

## Discussion

Our reconstruction of human germ cell lineages and their relationship to the somatic cells of the parents provides insights into the differentiation of early embryonic cells into germ and somatic cells. More specifically, the parental somatic presence of a germ cell mutation dates its emergence prior to PGCS and provides information about the population of cells selected for PGCS. The several cell lineages present both in the germline and soma, demonstrate that there are several cells contributing to the founding primordial germ cell population.

The variability between families in the contribution of these cell progenitors to the soma and germ cells, implies a stochastic sampling of cells to the germ and soma pools. Perhaps this highly variable sampling could be a byproduct of the spatial location of the cell lineages in the embryo, where related cells are selected as a group for PGCS. This scenario is supported by research of germ cells in mice, where cross signaling between cell layers determines a niche of cells that will become primordial germ cells^1^. Therefore, caution is warranted in the interpretation of simplistic tree-like modeling of the relative contributions from DNMs occurring in early cell divisions to observed somatic variation^5^. Interestingly, our data suggest that germ cells largely fall into two divergent lineages. A possible explanation for this is that the primitive streak formation at gastrulation, i.e. determination of the left and right symmetry in the embryo, dichotomizes the germ cell population into two parts that will subsequently populate the gonads.

Somatic variation is present in all tissue types of the human body. Although our sampling targets only a single tissue per proband (buccal or blood) this cell lineage sharing is expected to be present in other somatic tissues as the PGCS probably occurs at the onset of gastrulation^1^. Further, the number of ssDNMs observed in the large families indicates a significant accumulation of mutations in the divisions prior to the PGCS.

The filtering of germline DNMs with their presence in the somatic cells of parents biases estimates of mutation rates down, as mutations that are mosaic in the parents are generally omitted from mutation rate estimates. It has been proposed^13^ that this may make a significant contribution to the disparity between mutation rate estimated using fossil calibration in phylogenetic data (∼10^-9^ bp^-1^year^-1^)^25^ and the one using trios (4.27·10^-10^ bp^-1^year^-1^)^21^. However, this is not the case in our data as the sibling approach only increases the number of DNMs by 5%. We note that, while our sibling approach represents an improvement in identification of germline DNMs that are mosaic in parents, the requirement of at least one non-carrier with the same haplotype background means that we still miss DNMs that are present in all of the germ cells.

The DNM recurrence calculator presented here provides a tool for estimating the probability of recurrence of disease causing mutations in the clinical setting and allows parents to be better advised on future pregnancies.

## Methods

### Identification of ssDNMs in the trio approach

We used our previously described DNM set^21,26^. Briefly, to construct this DNM set we used DNM segregation in three generations families and quality covariates from GATK to construct a model to predict transmission of a DNM. Subsequently, we predicted proper segregation for the entire DNM set and the prediction was used to define high quality DNMs. We estimated the false positive rate in the DNM set to be low (2.9%) by the discordance among monozygotic twins in the set^21,26^.

We scanned this set of high quality DNMs for ssDNMs. Specifically, for a carrier of high quality DNM we determined whether siblings of the carrier also carried the DNM allele. We determined a sibling as a ssDNM carrier if there are at least two reads supporting the DNM allele, the allelic balance is strictly greater than 0.1 and the depth at the position is greater or equal to 10.

### Identification of early DNMs using siblings

We also identified DNMs by segregation in sibships where their genotype differs although they share the same parental haplotypes at a locus (Fig. 1b). For example, if a variant in an offspring is determined to be on a grand-paternal haplotype, then non-carrier siblings sharing the grand-parental haplotype at this position make the variant a DNM candidate.

We determine grandparental origin of haplotypes in the autosomal genome of the offspring based on sharing with the haplotypes of the parent. A haplotype shared with the parent on the maternal homolog of the parent is of grand-maternal origin and a haplotype shared on the paternal homolog is of grand-paternal origin.

We then scanned for DNM candidates using the grand-parental origin information for each sibship. In each sibship, we used carriers (defined below) to create a set of possible grand-parental origins of variants satisfying our criteria. That is, for each carrier we tabulated the possible grand-parental origins of the haplotype segments at the position of the mutation. We then calculated the intersection of the haplotype segments of the carriers to determine the set of possible grand-parental origins. We then tabulated whether there are siblings with the same set of possible grand-parental origins but that are non-carriers. If the carrier and non-carrier siblings share a grand-parental haplotype, we determined the variant to be a DNM candidate.

We defined a sibling as a carrier if the following criteria hold:

- minimum depth of 12 reads for the sibling
- minimum allelic balance of 0.15 for a carrier

We defined a sibling as a non-carrier if the following criteria hold:

- minimum 12 reads in the sibling supporting the reference allele
- maximum one read in the sibling supporting the alternative allele

To be confident in the grandparental origin of the haplotype, we filtered out siblings in the sibling approach if there is a cross over in the paternal or maternal chromosome within 500,000 bases. After extracting the set of DNM candidates via the sibling approach we validated the grandparental origin of the haplotypes by calculating the haplotype sharing between the carriers and the surrogate parents. To remove spurious DNM calls because of missed crossovers at the end of chromosomes we omitted DNM candidates less than 5Mb from the chromosomal ends. We only considered DNM candidates from the sibling approach if at least one of the carriers was present among the probands from the high quality DNM set derived from trio approach.

After this initial filtering we determined high quality DNMs using the same quality covariates as for the trios DNMs using the aggregated columns across the carriers. For example, we calculated the allelic balance across the carriers rather than individual carrier. These quality covariates were used as input into a generalized additive model described in^21^, which uses segregation of DNMs in three generation families to determine high quality DNMs. We annotated a DNM from the sibling approach not to be found if it was not among the trio DNM candidates.

We further excluded SNP DNMs from the ssDNM analysis that were less than 10 bases from an indel in the proband and all DNMs with a ratio of local coverage to the genome-wide average strictly greater than 1.8 or strictly less than 0.2.

#### Deep-sequencing parents of the large families

We deep-sequenced the parents of the 13 largest sibships to quantify the levels of somatic mosaicism in the parents and to ensure that the ssDNMs were not misclassified variants due to miscalled heterozygous genotypes in the parents. We added 3 HiSeqX lanes to the previous described WGS set^21^, resulting in 4 lanes per deep-sequenced individual or an average genome wide coverage of ∼172. The majority of the sibling pairs are from the large families (Supplementary Table 1), therefore even though only 13 of the 251 couples were deep-sequenced a substantial fraction of the sibling pairs has deep-sequenced parents (518 out of 1,007). For all13 deep-sequenced couples, DNA was extracted from blood. We applied the same sequencing and bioinformatics procedure as in the initial sequencing of the samples^26^, resulting in 26 merged BAM files.

We recalculated the parental allelic balance for all the DNMs in the set and estimated the background error rate by using the individuals in the WGS set which are not direct descendants of the couple. This was implemented with a custom python script using the pysam module (https://github.com/pysam-developers/pysam), we only considered aligned reads satisfying the following criteria:

- alignment of the read is primary.
- alignment is not a marked duplicate
- aligned read pair is a proper pair
- read is not flagged with a QC failure
- base quality of the read at the position of the DNM is greater than 17
- mapping quality greater than or equal to 10

For indels, we calculated the length difference between reference and alternative allele and compared that to the read alignment. We used the indel method of the pysam.PileupRead class to calculate the indel length supported by the read alignment and matched that to the DNM reference and the alternative allele length difference. The reads matching the allele length difference, were counted as supporting the alternative allele, and otherwise supporting the reference allele.

### Targeted resequencing of DNMs from the 17 and 10 sibling families

For the ssDNMs in the 10- and 17-sibling families, we PCR amplified in triplicates, a 1000 bp region around the DNMs in Supplementary Table 2. We ran each PCR on agarose gel and used the band intensity to normalize input amount (from 2-10 ul) of the PCR in 3 pools of all fragments for the two families. We fragmented the resulting 6 pools using a covaris E220 fragmenter and sequenced to a high sequence depth on a MiSeq instrument. The mean target size of the fragments was 300bp. We generated sequencing libraries using lllumina's TruSeq PCR-free sample preparation kit (FC-121-3003). We performed end repair, 3’-adenylation and ligation of indexed (96 dual indices) sequencing adaptors containing a T nucleotide overhang, followed by AMPure purification. We assessed the quality and concentration of all sequencing libraries using the LabChip GX instrument from Perkin Elmer. We pooled sequencing libraries (96 samples/pool) and sequenced on a MiSeq using the Illumina MiSeq v2 reagent kit (MS-102-2002). We performed the paired-end sequencing using 150 cycles for the forward and reverse reads respectively, in addition to the dual index reads. We demultiplexed and generated FASTQ files using MiSeq Reporter.

We aligned the reads in the FASTQ files with BWA mem^27^ (0.7.10-r789) and used the same reference as for the alignment of the whole-genome sequenced samples. We sorted the aligned reads with samtools^28^ and marked PCR-duplicates with Picard Tools (version 1.117). For most of the DNM sites (46/51) the targeted enrichment resulted in more than 500x coverage for 10 or more family members. Per DNM site in Supplementary Table 2, we extracted all variants from the whole-genome variant set in 400 bases from the DNM position. Subsequently, we constructed a variant graph per DNM site in Supplementary Table 2 with Graphtyper (‘graph’ command). We then called the genotypes of all of the individuals in both families using Graphyper at the DNM sites (‘call’ command). Finally, we compared the genotypes and allelic balance from the targeted resequencing and the whole-genome sequencing (Supplementary Fig. 4). We restricted the AB comparison to positions with over 500x coverage.

### Statistical analysis of sibling pairs

To allow comparison between the families of different sizes, we decomposed the families into pairs of siblings resulting in DNM and sibling pair combinations. There are phylogenetic relationships of the sibling zygotes within the family, thus there is possible intra family correlation when considering ssDNMs. In contrast, for offspring from different families to share a DNM a recurrence of a DNM is needed, therefore, independence between families is a reasonable assumption. In other words, there is a block-like covariance structure in the data and in order to accurately estimate the standard deviation of estimates and p-values, we applied a jackknife procedure leaving one family out at time. To compensate for the differential contribution of pairs from the different families, we weighted the pseudo replicates using the number of pairs from each family as weights. We used the formulas from ^30^ for the jackknife estimator and estimate of the standard deviation.

### Mutation class enrichment

We tabulated the DNM and sibling pair combinations according to whether they are ssDNMs or not for each mutation class. This results in a 2x2 table for each mutational class.

We calculated the odds ratio using the fisher.test function in R. We jackknifed the log odds ratios over the DNM and sibling pair combinations using the method described in “Statistical analysis of sibling pairs” to estimate the standard deviation of the log-odds ratio. We subsequently calculated the confidence interval for the log odds ratio using a normal approximation and transformed to the confidence interval for the odds ratio using the exponential function. We applied a similar procedure for phased ssDNMs, but instead of tabulating by ssDNMs status we tabulated DNM and sibling pair combinations according to phase and mutational class.

#### Determining DNM phase

DNMs determined by the sibling approach were phased using two previously described methods^21,26^, namely, three generation phasing and physical read tracing. Briefly, for the former if any of the DNM carriers had an offspring we phase the DNM by the transmission pattern of the DNM allele. For the latter, we physically read trace the DNM allele to nearby phased genotypes.

In addition to these approaches, we phased ssDNMs by haplotype sharing between DNM carriers. More specifically, if all of the carriers share haplotype of specific grand-parental origin at the DNM locus, we determine the parent-of-origin of the DNM depending whether the shared haplotype segment at the DNM locus is of paternal or maternal origin.

#### Phylogenetic reconstruction of cell lineages

We built a phylogenetic tree of the early cell lineages for each of the large sibships using the haplotype sharing between siblings and the set of ssDNMs. First, for all DNM sites we encoded the DNM presence at each locus across the siblings with three values: absent, carrier or missing. For ssDNMs of paternal origin, we determined a DNM to be absent from a sibling if the sibling was not a carrier and had the same paternal haplotype background as the DNM carriers at the DNM loci, but missing from the sibling if the sibling had the other paternal haplotype at the DNM locus. We used the corresponding definition for the maternal ssDNMs.

This resulted in a matrix of DNM sharing between siblings. To root the phylogenetic trees after the maximum likelihood estimation, an individual consisting only of absent calls was added to the matrix. Subsequently, this augmented DNM sharing matrix was used as input to RAxML^31^ (version 8.2.9) to estimate the phylogenetic tree of the cell lineages. As we only provide RAxML absence, carrier and missing status at variant sites, we used the RAxML model “-m ASC_BINGAMMA” and used the option “—asc-corr=lewis”. Raxml integrates over the possible genotypes at missing cells in the DNM sharing matrix, which is important, as half of the dataset is missing due to the haplotype background of the DNM is on average only observed in half of the siblings. The phylogenetic trees were constructed per family and parent-of-origin.

We used the aggregated DNM set from the trio and the sibling approach for this phylogenetic analysis, where we restricted to phased ssDNMs from the deep-sequenced families. In addition to the set of ssDNM, we incorporated DNMs that are somatic mosaic in the parent, i.e. >3 reads supporting the alternative allele in the parent.

The DNM sites in Fig. 2 and Supplementary Fig. 1 and 2 were sorted according to the following steps. First, we determined groups of DNM sites which have shared DNM carriers. Second, we calculated the fraction of carriers among carriers and absent. We then sorted the DNM sites by the group membership and then by the carrier fraction.

#### Pairwise dating of DNMs

In addition to the phylogenetic reconstruction we dated relatively all the pairs of DNM sites using the matrices described in the previous section. We used all of the DNM sites in the deep-sequenced families, i.e. corresponding to DNMs that are shared and not shared between siblings. For each DNM site combination (DNM site A and DNM site B) in the family, we tabulated a four way statistic using the carrier status of the siblings:

- F_00_: Sibling S1 and Sibling S2 are not carriers (absent, but not missing) of DNM site A and DNM site B
- F_10_: Sibling S1 is a carrier of DNM site A and Sibling S2 is not a carrier of DNM site B
- F_01_ Sibling S1 is not a carrier of DNM site A and Sibling S2 is a carrier of DNM site B
- F_11_ Sibling S1 and Sibling S2 are carriers of DNM site A and DNM site B

We restricted to pairs of DNM sites where F_11_>0 and either F_10_> 0 or F_01_>0, but not both. If F_10_>0 then we determine that DNM site B is relatively older than A (Fig. 3c). The reciprocal applies if F_01_>0 i.e. DNM site B is relatively younger than A. For each DNM site we calculated the average parental AB difference of the DNM site to all of its relative younger DNM sites.

### DNM recurrence calculator

The DNM recurrence calculator described in this section is available at

http://de-novo-risk.decode.is

using the following credentials

User:tester

Password:tester.pass!

We incorporated several covariates into the modeling of DNM recurrence, where the following covariates are of relative high importance: parental somatic mosaicism, presence in an older sibling and DNM phase. However, these covariates are not always available in clinical cases, thus our model accounts for the uncertainty or absence of these covariates.

In this section we use the following notation for observed values and parameters.

- S_y_, Boolean whether the DNM is shared by an younger sibling
- S_0_, Boolean whether the DNM is shared by an older sibling
- X_phase_, the phase of the DNM (“paternal” or “maternal”)
- *ω*, the vector of parameters

And the following notation for covariates

- A_p_, age of father at conception
- A_m_, age of mother at conception
- C, mutation class of the DNM
- *M_P_*, paternal somatic presence (SMPP described below)
- *M_M_*, maternal somatic presence (SMPP described below)
- *G*, Boolean whether the DNM is within a C>G enriched region

In the DNM recurrence modeling we considered trios of siblings rather than pairs of sibling and integrate out the older sibling occurrence when information from multiple siblings is not available. For the training of the recurrence calculator we only considered trios of siblings where both of the parents the DNA was extracted from blood.

#### Somatic mosaicism

Accurate determination of levels of somatic mosaicism can be time and cost prohibitive for clinicians as high sequencing depth is needed. Further, half of the sibling pairs in our data set do not have parents of high sequence coverage. With these considerations in mind, we devised a statistical framework taking into account the uncertainty of the somatic mosaic estimation into the recurrence modeling.

We used the read counts of the deep-sequenced parents across all DNM sites to construct an empirical prior distribution of the parental somatic mosaicism. The prior distribution calculation was restricted to the set of deep-sequenced parents of the 13 largest sibships as they have the most accurate characterization of the somatic mosaicism and have the greatest power to detect DNMs filtered due to parental presence.

We fitted a beta-binomial distribution to the empirical read counts supporting the reference and alternative alleles at all DNM sites for both parents. The fitting was implemented with a likelihood approach, using the optimization function optim in R using the “BFGS” optimization method. To avoid local optimal likelihood values due to badly selected starting values, we started the optimization in all possible α and β combinations of the following values 0.01, 0.1, 1,10 and 100. The parameters corresponding to the maximum likelihood value over all of the starting values was used in the subsequent analysis. The probabilities of the beta binomial distribution were calculated with the dbb function from the TailRank R package.

For all DNM and sibling trio combinations, we calculated the posterior distribution of the somatic mosaicism of the parent using the fitted beta distribution as a prior. From this posterior distribution, we made a new covariate (somatic mosaicism posterior probability; SMPP) by calculating the posterior probability of observing allelic balance greater than 1%. We removed sites with high SMPP rates for both parents (PSMPP>0.8). Further, we aggregated the reads counts for all individuals that are not descendants of the parent pair (background) and calculated the PSMPP rate for the background. We filtered out DNM sites where the background had SMPP rate of 0.8 or greater.

#### Predicting the parent-of-origin of unphased DNMs

Determination of the parent-of-origin of DNMs is often impossible, thus we incorporated the phase uncertainty into the calculation. More specifically, we predicted the parent-of-origin of the DNMs using the following attributes in a generalized additive model formula (GAM):

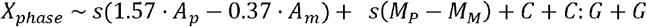

where 1.57 and 0.37 are our estimates of the increase in DNMs with parental and maternal age, respectively^21^. The sibling trios and DNM combinations were weighted in the GAM fitting such the contribution of each DNM site to the likelihood was equal.

#### The recurrence probability likelihood

The likelihood for the recurrence part of the model is the following

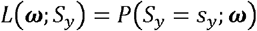

We sum out the phase of the DNM and the older sibling status.

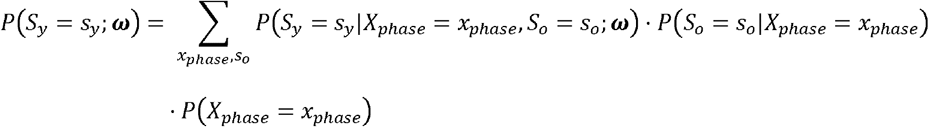

For phased DNMs, *P*(*X_phase_ = x_phase_*) was either 0 or 1, depending on parent-of-origin, and for unphased DNMs the estimated phase probability from the GAM was used. We used sibling trios for the likelihood, thus *P*(*S_o_* = *s_o_*|*X_phase_* = *x_vhase_*) was set to 1 or 0 depending on whether the older sibling was a DNM carrier or not, respectively.

We then model sex-specific probabilities that the DNM is shared by with a logistic link function, i.e.

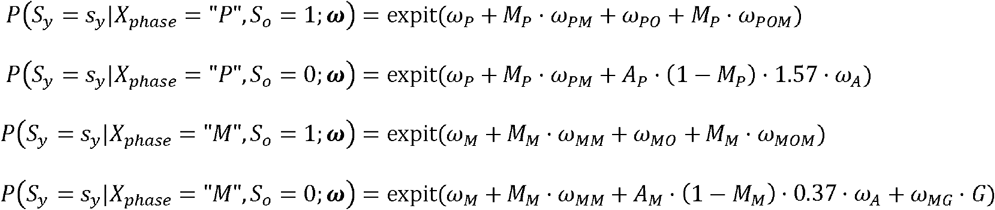

The absence of the age covariates in the case of S_o_ = 1 is to capture that multiple transmissions are informative about the timing of DNM, i.e. the DNM probably occurred early in the embryogenesis of the parents.

Then for all trio combinations we calculate the likelihood

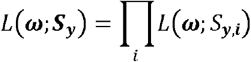

We estimated the maximum likelihood of the parameters using the BFGS method in the optim function in R. The initial BFGS iteration was started in this set of parameters:

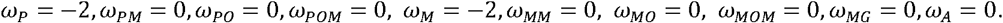

#### Summing out the older sibling status

When the carrier status of an older sibling is not available we predict the recurrence probability using the weighted average of the predictions based on the older sibling being a carrier or not:

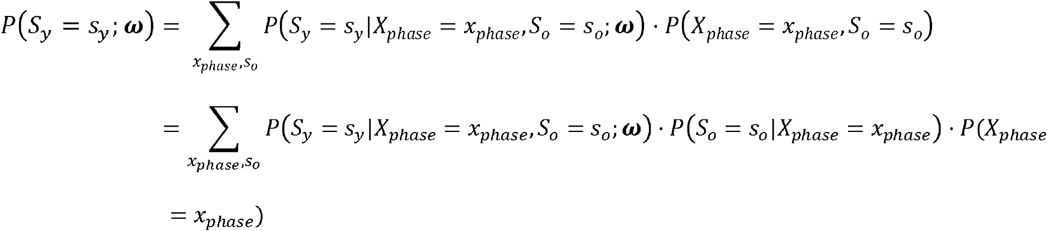

For *P*(*S_o_* = *s_o_*|*X_phase_* = *x_vhase_*), we use the empirical frequencies observed in the dataset of phased DNMs.

### STXBP1 clinical case

We were presented with a clinical case of two siblings with recurrent seizures, both of whom were under 5 years of age at the time (Fig. 5b). Further, their parents do not have any history of seizures. To determine whether the affected siblings’ condition had a genetic origin, we whole-genome sequenced the siblings and the unaffected parents and scanned for rare variants described or predicted to be pathogenic.

DNA isolated from blood samples from the siblings and their parents was prepared for whole-genome sequencing following the TruSeq Nano sample preparation method. The samples were sequenced on Illumina HiSeqX machines to a targeted depth of 30X. Sequence reads were aligned to NCBI’s Build 38 of the human reference sequence using version 0.7.10 of the Burrows-Wheeler Aligner^27^ and were merged into BAM files. Reads were marked for duplicates using Picard 1.117, only non-duplicate reads were used for the downstream analyses.

Variants were called with version 2.3-9 of the Genome Analysis Toolkit^32^ using joint calling with HaplotypeCaller, version 2014.4-3.3.0-0-ga3711aa. We restricted the subsequent analysis to SNPs and small indels (shorter than 20 base pairs) at coding and splicing regions, as annotated by release 80 of the Variant Effect Predictor^33^ using RefSeq gene annotations. We focused on rare genotypes in genes where pathogenic mutations are known to contribute to Mendelian disease as annotated in OMIM^34^. We counted the number of reads supporting the reference and alternative allele using pysam (version 0.8.3, quality_threshold=15 and read_callback=‘all’).

We defined rare autosomal recessive genotypes as homozygous or compound heterozygous sequence variants, each with a minor allele frequency lower than 1% in our set of 15,220 whole-genome sequenced individuals^21,26^ and gnomAD^35^. We defined rare autosomal dominant genotypes as variants with a minor allele frequency lower than 0.1% in our set and gnomAD.

We detected no rare homozygous or compound heterozygous genotypes shared by the two affected siblings at coding/splicing regions in their genomes. We detected no rare, heterozygous genotypes shared by the siblings that were absent from their parents. Out of 148 moderate or high impact variants (coding/splicing) with an allelic frequency under 0.1% present in both siblings (and in either parent), we observed 78 variants in previously described disease genes (OMIM). Among these we found a heterozygous missense variant, NP_003156.1:p.Gly543Glu (chr9:127,682,486 G>A), in the known epileptic encephalopathy gene *STXBP1*^23’24^. None of the other 77 disease genes had a reported link to a seizure phenotype (OMIM, Human Phenotype Ontology^36^). The *STXBP1* mutation, shared by the siblings and their father (Fig. 5b, Supplementary Table 3), is absent from the mother, our set of 15,220 whole-genome sequenced individuals^21,26^ and gnomAD. The mutation is predicted to be deleterious by *in silico* tools SIFT^37^ and PolyPhen-2^38^. Further, the chromosomal position of the mutation, chr9:127,682,486 (GRCh38), is highly conserved evolutionarily (GERP^39^ score of 5.62).

We saw evidence for mosaicism of the mutation in the father’s blood sample (AB=12%; p=0.00016 binomial test of 50%, Supplementary Table 3). The paternal somatic mosaicism was verified with bidirectional Sanger sequencing of both his blood sample and a new buccal sample (Supplementary Table 3). Sanger sequencing confirmed heterozygous status of the mutation in the two affected siblings, as well as the absence of the mutation from their paternal grandparents (buccal tissues). The mosaic presence in the father’s soma and the transmission of the allele to multiple offspring indicates that the mutation is a DNM that occurred before the PGCS in the father.

*STXBP1* missense mutations are an established cause of early infantile epileptic encephalopathy (OMIM) and in fact, paternal mosaicism of a pathogenic *STXBP1* mutation has been reported before^40^. No other likely pathogenic variants were found during our clinical analysis of the genomes of the affected siblings. The absence of the *STXBP1* DNM from Icelandic controls (15,220 whole genome sequenced individuals) and from gnomAD, as well as the mosaic presentation of the DNM in the father strongly indicates that this is the causative genotype.

#### Genotype validation using the Sanger sequencing method

We designed primers with the Primer 3 software^41^. We performed PCR and cycle sequencing reactions on MJ Research PTC-225 thermal cyclers, using the BigDye Terminator Cycle Sequencing Kit v3.1 (Life Technologies) and Ampure XP and CleanSeq kits (Agencourt) for cleanup of the PCR products. We loaded sequencing products onto the 3730 XL DNA Analyzer (Applied Biosystems) and analyzed with the Sequencher 5.0 software (GeneCodes Corporation). We estimated the relative peak heights for each individual with the Sequence scanner software 2.0 (Applied Biosystems Inc/ThermoFisher Scientific).

